# Decoding neurobiological spike trains using recurrent neural networks; a case study with electrophysiological auditory cortex recordings

**DOI:** 10.1101/2020.12.14.422705

**Authors:** Péter Szabó, Péter Barthó

**Affiliations:** Institute of Evolution, Centre for Ecological Research, H-1121 Budapest, Hungary; MTA TTK NAP B Sleep Oscillations Research Group, H-1117 Budapest, Hungary

**Keywords:** neural code, auditory cortex, population coding, recurrent network

## Abstract

Recent advancements in multielectrode methods and spike-sorting algorithms enable the in vivo recording of the activities of many neurons at a high temporal resolution. These datasets offer new opportunities in the investigation of the biological neural code, including the direct testing of specific coding hypotheses, but they also reveal the limitations of present decoder algorithms. Classical methods rely on a manual feature extraction step, resulting in a feature vector, like the firing rates of an ensemble of neurons. In this paper, we present a recurrent neural-network based decoder and evaluate its performance on experimental and artificial datasets. The experimental datasets were obtained by recording the auditory cortical responses of rats exposed to sound stimuli, while the artificial datasets represent preset encoding schemes. The task of the decoder was to classify the action potential timeseries according to the corresponding sound stimuli. It is illustrated that, depending on the coding scheme, the performance of the recurrent-network based decoder can exceed the performance of the classical methods. We also show how randomized copies of the training datasets can be used to reveal the role of candidate spike-train features. We conclude that artificial neural network decoders can be a useful alternative to classical population vector based techniques in studies of the biological neural code.

## 1 Introduction

Nerve cells use action potentials (AP), rapid electrical impulses, to transmit information. The question how the temporal sequences of these action potentials encode the biologically relevant information is a major, unresolved issue in neurobiology. According to the rate encoding hypothesis, which dates back to Adrian and Zotterman [1], the information is represented by the number of spikes within an appropriate time window, irrespective of their temporal distribution. However, both experimental results and theoretical considerations suggest that the complexity of the neural code goes beyond this simple scheme [2,3]. The temporal coding hypothesis emphasizes the role of the precise timing of action potentials. The wide range of candidate hypotheses involve diverse AP timing features, like latencies, relative phases, correlations or synchrony. Another line of extension is population encoding; the information is encoded in the combinations of the different firing neurons. A demonstrative example is rank order coding, were encoding is based on the temporal order of firing cells [4].

One approach for the study of neural code is to compare the expected biological performance of the different encoding schemes from an information-theoretic viewpoint [5–8]. However, since current electrophysiological methods are able to record AP sequences simulatenously from many neurons with high temporal resolution, it is also possible to assess the biological relevance of the encoding schemes directly [9–11]. The experimental datasets can be used to fit models that reconstruct the signal from the emitted AP timeseries using some specific encoding scheme. Most of these studies use population vector representation of the AP timeseries. These vectors contain the firing rates of the different cells, i.e. they integrate the features of rate and population encoding. Population vectors are easy to construct and they perform well in many cases, still, the analysis involves a manual feature extraction step, which constrains the range of involved features.

The demonstrated ability of artificial neural networks to function as general function approximators, i.e. their capability to learn the functional relationship between interrelated variables, makes them promising tools for studying neural coding, because the objective is to find a mapping from the action potential sequence to the encoded message. Since artificial neural networks extract the relevant features during their training, the input data can be the raw action potential timeseries, avoiding any constraint on the involved features. Recurrent neural networks (RNN), and mostly their more advanced long short-term memory (LSTM) and gated recurrent unit (GRU) variants, are especially adept at analysing sequential information due to their internal memory state, that keeps track of the information that is gradually extracted from subsequent elements of a sequence. The main application area of RNNs is natural language processing (NLP), they are integral parts of machine translation algorithms, but they are already extensively used also in several fields of biology, including genomics [12,13], proteomics [14,15] and neurobiology [16,17]. Their popularity in these fields is partly due to the existence of reliable, large databases which are the most important prerequisites of any machine learning methods.

Our approach of applying RNNs for the study of neural encoding is based on the premise that neural action potential timeseries can be considered as linguistic sentences, where action potentials of different cells correspond to different words. Just as the meaning of sentences is determined by the pattern of words in the sentence, a neural “message” is determined by the temporal pattern of action potentials of neural cell ensembles. This is similar to the NLP interpretation of the relationship between DNA and protein sequences or the relationship between the structure and function of enzimes. The issue of neural decoding can be considered as a classification or regression task; one needs to assign the belonging stimulus to an action potential timeseries. In the followings we present a RNN neural network based decoder, which learns the encoding rule from action potential timeseries datasets. We use both experimental and artificial datasets to illustrate the performance of the RNN decoder. Since the learning process itself does not provide information about the acquired decoding rule, we complement the study with a perturbation analysis to reveal the role of different features in the signal representation.

## 2 Materials and methods

We used six datasets (summarized in Table 1). The AP timeseries are represented as *L* = 500 long vectors of integers, each belonging to one of *S* = 18 stimulus identifiers. The vector elements express the occurrences of action potentials in subsequent 1ms long bins following the onset of the stimulus; zero values indicate that no action potential occurred in the given bins, whereas positive integers identify the firing neurons. The three experimental datasets (D^E^_I–III_) were obtained by recording the auditory cortical responses of rats exposed to sound stimuli, while the three artificial datasets (D^A^_I–III_) were obtained by simulating preset encoding schemes.

**Table 1.** Summary of the action potential timeseries datasets. Superscripts ‘A’ and ‘E’ denote artificial and experimental datasets, respectively. ‘Items’ is the number of spike timeseries in the dataset, ‘Cells’ is the number of involved neurons.

| Dataset | Items | Cells | Notes |
| --- | --- | --- | --- |
| $D^E_I$ | 3618 | 51 | auditory cortex |
| $D^E_{II}$ | 4032 | 70 | auditory cortex |
| $D^E_{III}$ | 1458 | 70 | auditory cortex |
| $D^A_I$ | 1800 | 50 | frequency encoding |
| $D^A_{II}$ | 1800 | 37 | population encoding |
| $D^A_{III}$ | 1800 | 5 | rank order encoding |

### 2.1 Experimental datasets

The experimental datasets (D^E^_I–III_) were collected from auditory cortex recordings of four Sprague–Dawley rats (300–450 g), as described in [18]. Briefly, animals were anesthetized with urethane (1.5 g / kg) and placed in a custom-made naso-orbital stereotaxic apparatus. Recordings were made from the auditory cortex with 32-channel silicon microelectrodes (8 shank, tetrode configuration, Neuronexus Tech, Ann Arbor, MI, USA) at a depth of 1-1.5 mm. Electrodes were estimated to be in deep layers by field potential reversal [19], most likely layer V due to electrode depth and the presence of broadly tuned units of high background rate [20]. The location of the recording sites was estimated to be the primary auditory cortex (A1 / AAF) based on stereotaxic coordinates, vascular structure [21–23], tonotopic variation of frequency tuning across recording shanks, and the presence of cells with V-shaped tuning curves. Extra-cellular signals were band-pass filtered (1–8 kHz) and amplified (1000 times) using a 64-channel amplifier (Sensorium, Charlotte, VT, USA), and digitized at 20 kHz. Units were isolated by a semi-automatic algorithm [24], followed by manual adjustment (http://klusters.sourceforge.net). Multi-unit activity, clusters with low separation quality (isolation distance *<* 20) were excluded from the analysis [25,26]. All experiments were carried out in accordance with protocols approved by the Rutgers University Animal Care and Use Committee, and conformed to NIH Guidelines on the Care and Use of Laboratory Animals.

Experiments were conducted in a sound attenuating chamber. Sounds were generated by an RP2 signal processor, attenuated by a PA5 attenuator, and delivered free field by an ED1-ES1 speaker system (Tucker-Davis Technologies, Alachua, FL, USA). The stimulus battery consisted of 18 pure tones logarithmically spaced at 3–43 kHz. To compensate for the transfer function of the acoustic chamber, tone amplitudes were calibrated prior to the experiment using a condenser microphone placed next to the animal’s ear (7017, ACO Pacific, Belmont, CA, USA) and an MA3 microphone amplifier (Tucker-Davis). The stimuli and the subsequent silent intervals were 1 s long in datasets D^E^_I–II_ 0.5 s long in dataset D^E^_III_, at 70 dB SPL in all cases. Tones were presented repeatedly at 70 dB SPL in random order.

The action potential vectors were obtained by merging the spike trains of individual cells within subsequent 1 ms long bins (Fig. 1). If none of the identified cells fired during a bin, then the vector element was assigned a value of zero, otherwise it was assigned the ID of a randomly selected cell from among the firing neurons. Due to the sparseness of spikes (*≈* 0.1*/*ms), the spike loss, originating from the binning and merging operations, was minimal.

**Fig. 1.**
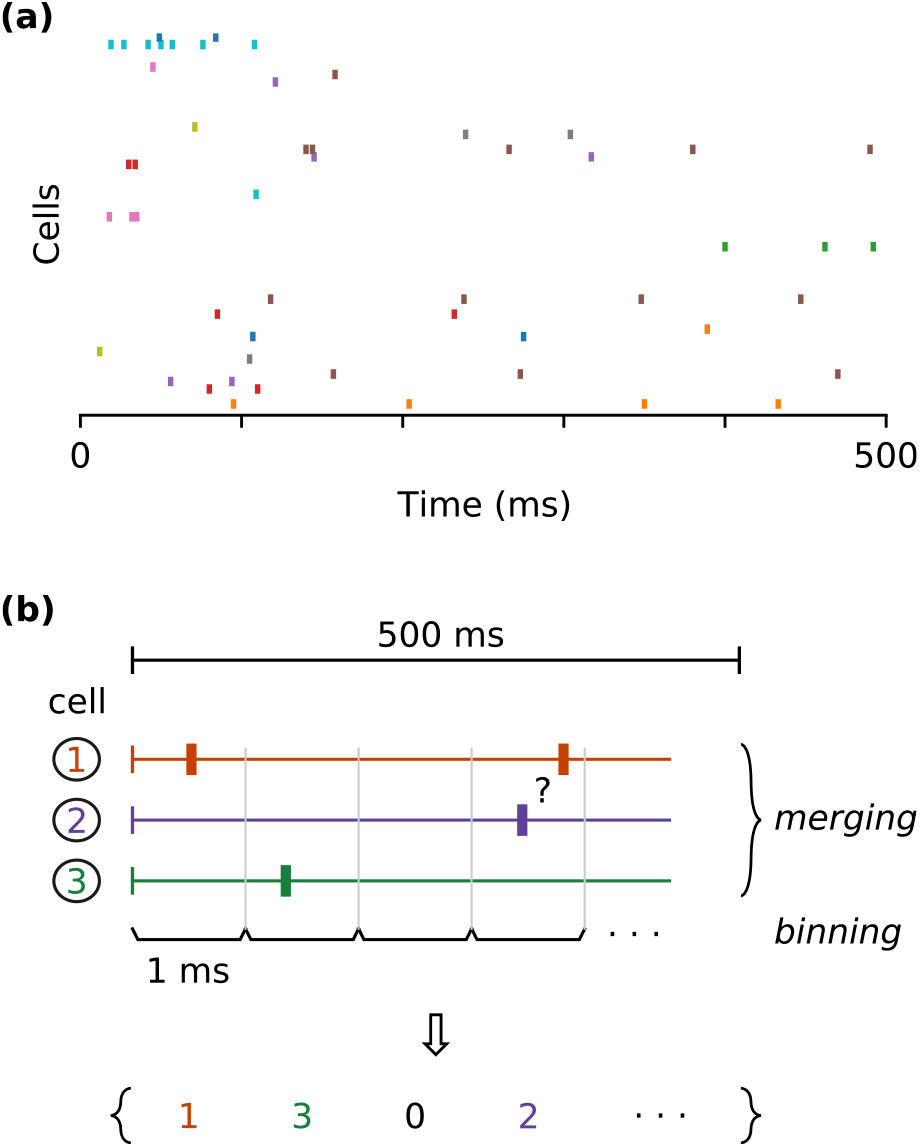
Preprocessing of the experimentally recorded cell-specific action potential timeseries. **a** A sample timeseries. The action potentials of different cells are denoted by unique colors. **b** Vectorization process; the timeseries of individual cells were merged into a single vector by assigning the identifiers of firing cells to subsequent 1 ms-long time bins. If none of the cells were activated within a time bin (bin 3 in the example), then the corresponding vector element was zero. If more then one cell emitted an action potential within the same time bin (bin 4 in the example), then the corresponding vector element was a randomly selected identifier from among the firing cells.

### 2.2 Artificial datasets

The artificial datasets were obtained by using simulated data with preset encoding schemes. In dataset D^A^_I_, the stimulus was encoded by the overall frequency of spikes (frequency encoding). An *i*^th^ element of a timeseries vector was nonzero with a stimulus-specific AP probability of 0.05 *· s*, where *s* stands for the stimulus identifier (*s* = 1, …, *S*). For each spike, the corresponding cell identifier was a random value from the range 1–50. In D^A^_II_, the stimulus was encoded by the ensemble of activated cells (population encoding). An *i*^th^ element of a timeseries was nonzero with a fixed spiking probability of 0.1, while the belonging cell identifiers were random values from the stimulus-specific cell triplets {2*s* − 1; 2*s*, 2*s* + 1}. In D^A^_III_, the stimulus was encoded by the order of the firing cells (rank order encoding). First, each stimulus was assigned to a unique permutation of five cell identifiers (excluding cyclic-invariant permutations). An *i*^th^ element of the timeseries was nonzero with a fixed probability of 0.1, and the belonging cell identifiers were (cyclically) consecutive elements of the stimulus-specific cell permutations. For each encoding scheme, we created 100 simulated timeseries for each stimulus.

### 2.3 RNN decoder

We used an artificial neural network decoder to receive an AP timeseries as input and give predicted likelihood scores for the possible stimuli as output. The decoder itself was a standard recurrent neural network with self-attention mechanism, with its structure shown in Fig. 2. The input is a one-hot encoded AP timeseries of shape (*L, C*), where *C* is the number of cells in the dataset. The decoder processes this data in three major steps: cell embedding, sequence embedding and classification. During the first, cell embedding step, the one-hot cell vectors are replaced by learnt vectorial cell representations of length *E* = 50, resulting in an array of shape (*L, E*). This is formally accomplished by multiplication with a cell embedding matrix, which is responsible for learning functional cell representations that express the role of the different cells in the stimulus decoding. The second, sequence embedding step is performed by a recurrent network enhanced with a self-attention mechanism. The recurrent network consists of a single layer with *H* = 400 GRU units. A GRU unit [27] is a gating mechanisms that enables the addition or removal of information from a memory thread. They are simpler and often more efficient than the alternative LSTM gating units. These units process the bins of the timeseries in a sequencial order, extracting relevant information during each step as hidden state ouputs. This hidden state information is then processed with a self-attention mechanism, which linearly combines the list of hidden-state outputs using context specific attention weights. The attention mechanism [28–30] helps the RNN to context dependently focus on certain parts of the input sequence, enabling easier and more efficient learning. This results in an embedded sequence representation matrix of shape (*D, H*), where *D* = 32 is a feature number parameter of the attention mechanism. The final, classification step is done by a two-layered fully connected network. The first layer consisted of 1000 cells with sigmoid activation functions, whereas the second, output layer had *S* number of cells with softmax activation funtions. The final classification layer provides likelihood scores for all the stimulus classes; the single predicted (encoded) stimulus is the one with the highest score. The trainable parameters of the model are the cell-embedding matrix, the GRU network weights, the attention matrix, and the fully connected network weights.

**Fig. 2.**
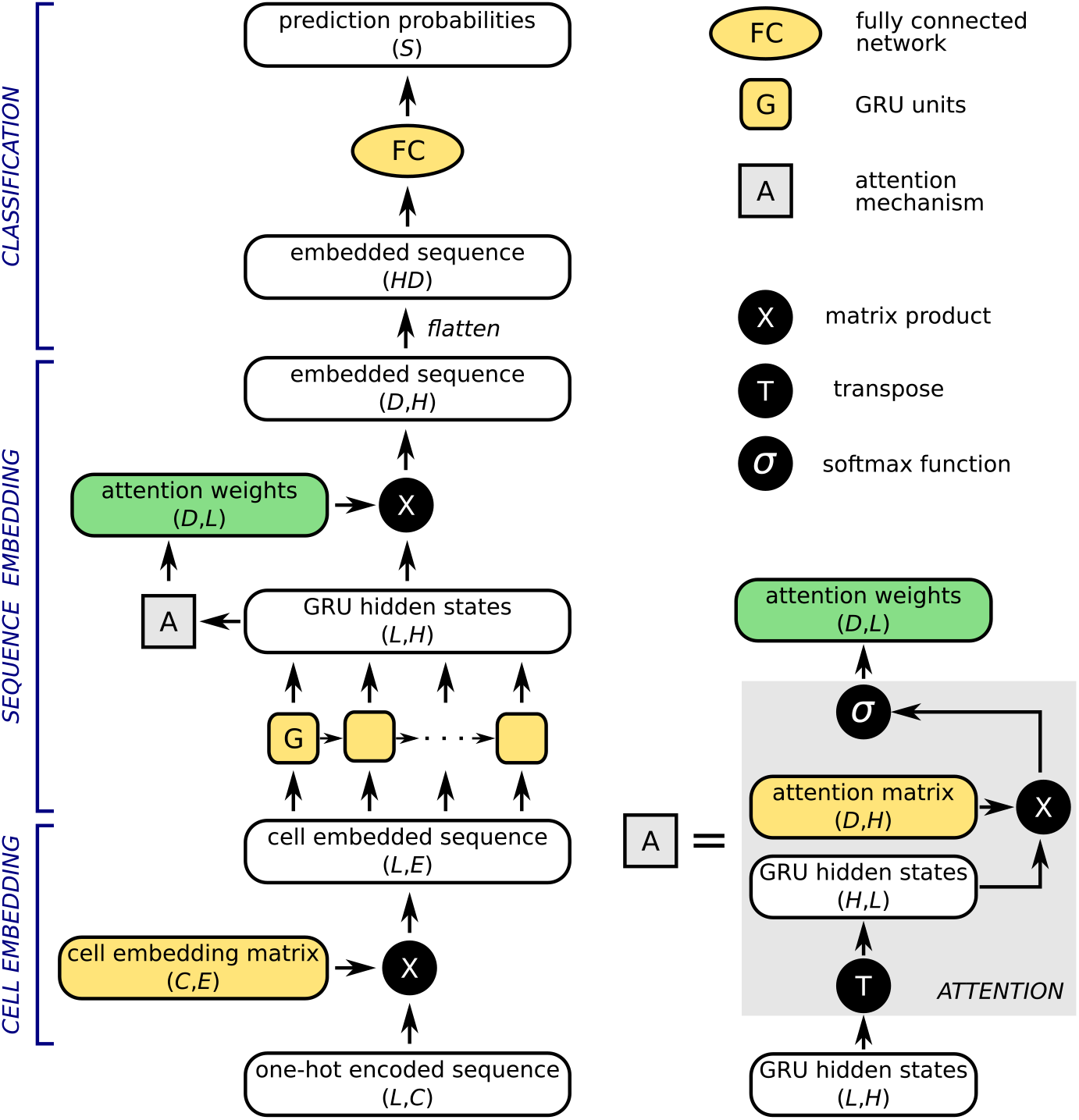
Structure and functioning of the recurrent neural network decoder. The network has three major components, which perform the cell embedding, sequence embedding and classification tasks. The sequence embedding part contains also a self-attention mechanism, which is depicted in details in the lower right panel. White boxes represent the data in different stages of the processing pipeline with their shapes shown in parentheses (the batch dimension is not displayed). The trainable parts of the network (cell-embedding matrix, attention matrix, weights of the GRU units and the fully connected network) are denoted by yellow. *L* denotes sequence length, *C* stands for the number of neurons in the dataset, and *S* is the number of stimuli. *H* stands for the number of GRU units, while *E* and *D* are feature number parameters in the cell embedding and attention matrices, respectively.

We used Adam optimizer with a cross entropy loss function and L2 (or ridge regression) regularization (learning rate=10^−4^, L2 weight decay=10^−5^, no. epochs=200, batch size=100). The L2 regularization helps to reduce overfitting by adding a penalty term to the loss function that forces the algorithm to keep the model weights small. Prior to analysis, each dataset was divided into training, validation and test sub-datasets in the ratio of 80:10:10. The training datasets were used to train the model during the epochs, while the validation datasets were used to prevent the model from overfitting (the model weights were saved at minimal loss for the validation dataset). We ran eight independent training sessions for each dataset with random initial weights, and the models with the lowest evaluation losses were used on the test datasets to assess the prediction accuracies. Fig S1 illustrates the training session on dataset D^E^_I_.

### 2.4 LDA decoder

For comparison, we also performed a more classical analysis of the same datasets using population vectors and linear discriminant analysis (LDA). The population vectors were calculated from the action potential timeseries by calculating the number of APs belonging to different cells within 50ms length bins. Note that this is a flattened vector, which contains the firing rates of all cells in all the time bins. Similar to the RNN decoder, the LDA model was fitted to the train datasets, whereas the prediction accuracy was assessed by applying the fitted model to the test datasets.

### 2.5 Perturbation analysis

In order to infer the role of different kind of information in the stimulus encoding, we compared the performance of the two decoders against the original test datasets with their performances against perturbed variants of the test datasets. The perturbations were meant to selectively remove certain types of information from the original data in the following way. Each dataset can be considered as a three dimensional array of size (*S, R, T*), where the values are organized along the stimulus, repeat and time axes. Shuffling the values within the vectors along any of these axes removes the axis specific information. If we shuffle the values along the repeat dimension (R perturbation, meaning the shuffling of values with the same *s* and *t* indices) then we remove any correlations within the AP timeseries bins; the probability of different spikes within each bin will be equal to the stimulus and time-specific average value. Shuffling the values along both the repeat and time dimensions (RT perturbation) retains only the stimulus-specific averages. Shuffling along all the there dimensions (RTS perturbation) results in a dataset where the occurrences of spikes and cells in all of the bins correspond to their global average probabilities.

Applying shuffles in the above way (called spike perturbations) implies the perturbation of both AP locations and cell identifiers, but we can also make shuffles affecting only the cell identifiers (cell perturbations), which essentially means a reassignment of the cells among the action potentials, while retaining the original chronology of spikes. Cell perturbations will be denoted by accented axis names (R′, T′*and*S′). Note that, since a spike perturbation always implies also a cell perturbation, the two types of perturbations can not be combined completely freely. E.g. an RS spike perturbation can be performed with or without a T′ cell perturbation, but an RST perturbation can not.

## 3 Results and Discussion

Fig 3 shows the main descriptive statistics of the experimental datasets. The auditory cortex responds to stimuli with transient discharges across a relatively large population of neurons. In all cases, the overall firing rates show a larger and smaller peak at around 25-50ms and 160ms, respectively, before approching the mean value after 250ms. The distribution of action potentials among the neurons is higly unequal; a small portion of cells is responsible for the majority of the spikes. (Fig S2 shows the same statistics for the artificial datasets.)

**Fig. 3.**
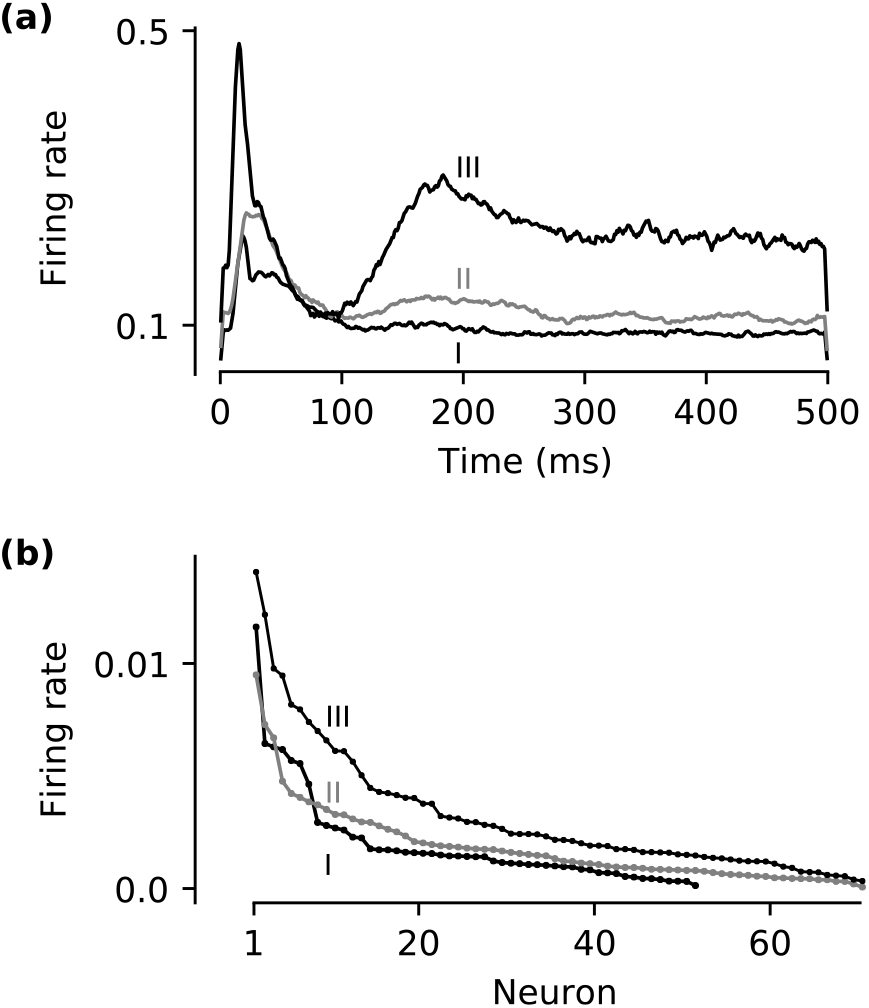
Descriptive statistics of the timebin- and cell-specific firing rates in the experimental datasets. **a** Overall firing rates, summed over all cells, in subsequent time bins after the stimulus onset. The timeseries were smoothed with a flat window, five bins wide. **b** Overall firing rates, averaged over time bins, of different neurons in decreasing order.

The stimulus information can be extracted from the timeseries with relatively high accuracy (Fig. 4). The RNN decoder classified the experimental sequences (D^E^_I–III_) correctly (i.e. predicted the correct stimulus) with accuracies 50, 54 and 43 percents, respectively. The corresponding values for the LDA decoder were 53, 55 and 42 percents. For artificial frequency encoding (D^A^_I_) the RNN decoder performed better (79% vs. 54%), for artificial population encoding (D^A^_II_) both methods were very accurate (100%). For the artificial rank order encoding (D^A^_III_) there was a sharp difference between the performances; in contrast to the perfect (100% accuracy) predictions of the RNN decoder, the LDA performed poorly (9% accuracy). Since both methods performed roughly equally well for the experimental datasets and for artificial frequency and population encoding data, and the population vectors used by the LDA extract exactly this kind of information, the experimental AP timeseries probably also involved some frequency and cell population encoding components.

**Fig. 4.**
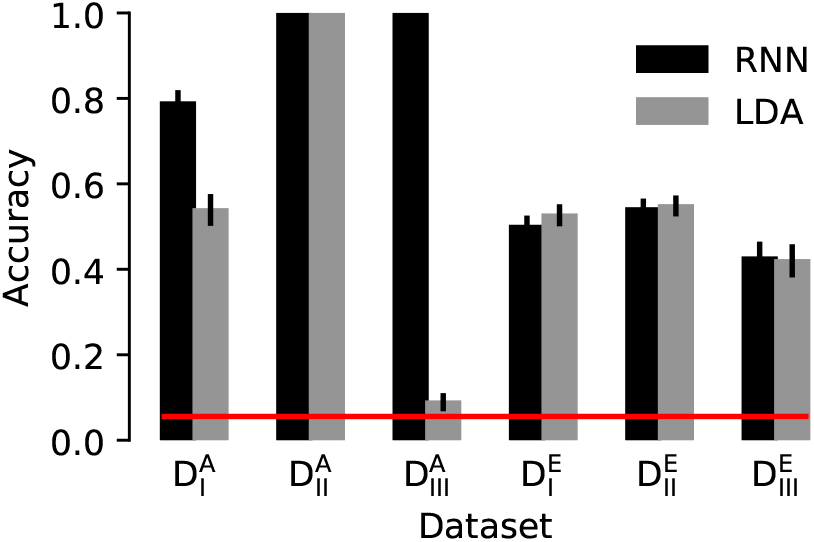
Prediction accuracies of the RNN and LD decoders. Accuracy is measured by the proportion of correct stimulus predictions in the test datasets. Error bars indicate the standard error. The red line shows the reference accuracy attainable by random choice (1*/S*).

The aim of the perturbation analysis (Fig. 5) was to reveal the role of different components by selectively removing some information from the original data, and comparing the decoding efficiencies for these perturbed data with those for the original one (see e.g. [31] for a similar approach). The use of this analysis is well illustrated by its application on the artificial datasets. Considering the frequency-encoded D^A^_I_ dataset the only perturbation that decreased (in fact completely ruined) the stimulus predictability is if we shuffle all values between all timeseries (RTS), which effectively means evenly redistributing the action potential events. Neither perturbation between repeats (R) nor perturbation between repeats and time-bins (RT) had any effect on predictability, since the timeseries contain no correlative information and the distribution of action potentials along the timebin axis is irrelevant. Likewise, cell perturbations did not decrease the predictability, because the identity of firing cells was irrelevant in this encoding scheme. In contrast, for the cell-population encoded D^A^_II_ dataset, the redistribution of the identity of firing cells between all timeseries (R′T′S′) was needed to decrease the predictability, because the timing of action potentials is now irrelevant. The order-encoded D^A^_III_ dataset contains correlative information, i.e. the “meaning” of a particular firing cell depends on the identity of previous and subsequent firing cells. It is reflected in the sensibility of the predicion power also against R/R′ type perturbations, which exhanged cell identities between repeats, completely wasting the cell order information within the individual sequences, even without a stimulus-wise (S) shuffle. Notice that, the detrimental effect of R′ cell-perturbations is smaller than R spike-perturbations, although they should be the same. This is because of the limited dataset size; in some time-bins only a single AP occurred among the different repeats; in these cases the cell-shuffles were futile.

**Fig. 5.**
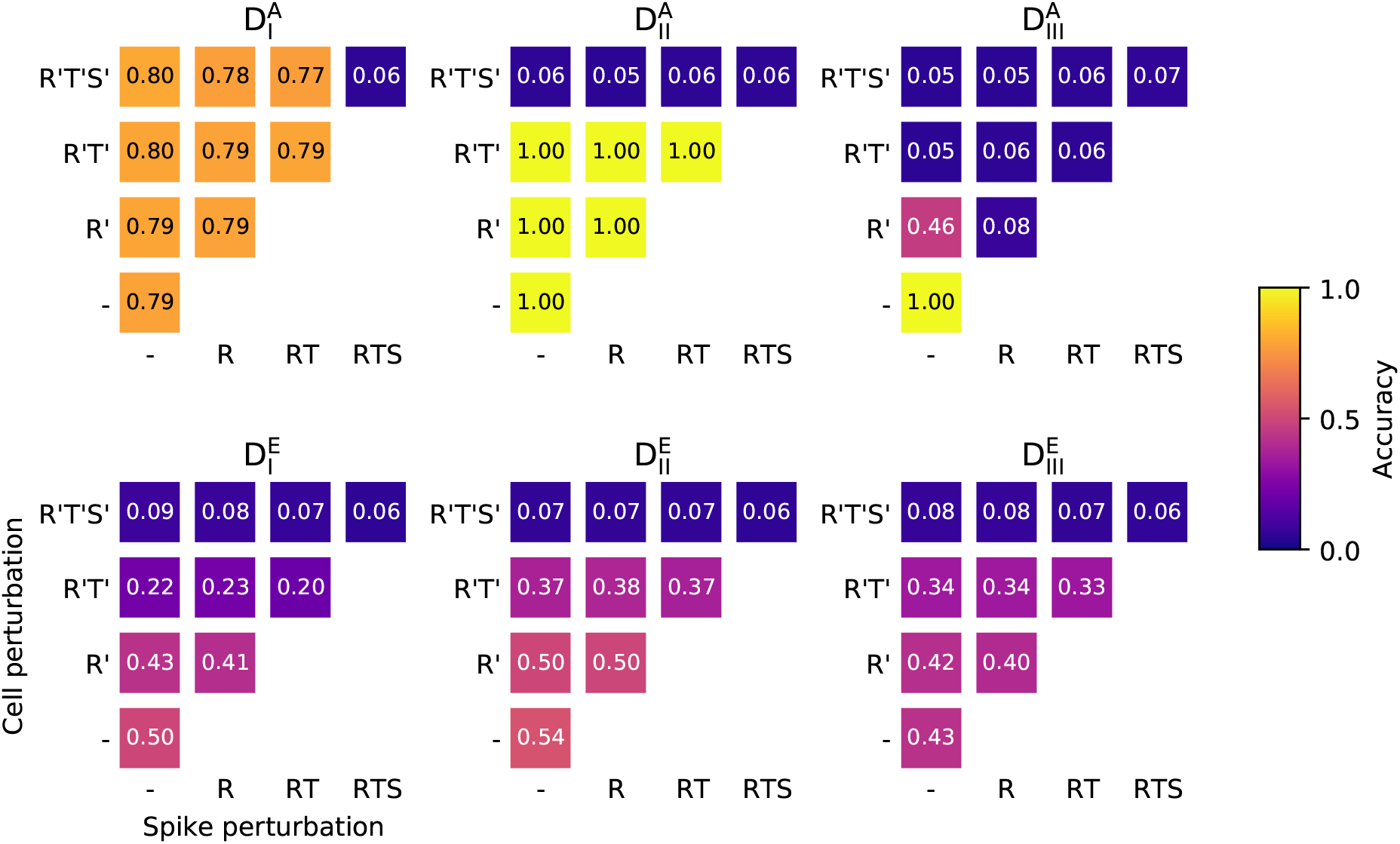
Perturbation analysis for the RNN decoder. The prediction accuracy of the RNN decoder for the original datasets is compared with the prediction accuracies for the perturbed datasets. Perturbations may affect either any of the values (spike perturbations) or only the positive values (cell perturbations) within the datasets. R, RT, and RTS spike perturbation names (and their primed counterparts for cell perturbations) refer to the affected repeat, time, and stimulus axis combinations (see main text for details). The standard errors of the accuracy estimations were less than 0.0123 in all cases.

In contrast to the deliberately simplified and sharply different artificial datasets, the experimental datasets exhibited a more intricate response towards various perturbations. Most importantly, only cell-perturbations had a strong negative effect on predictability. The negligible effect of AP location changes can be seen from the very similar accuracy values belonging to the same cell-, but different spike-perturbations. However, the comparison of the R′ perturbations with the R′T′ perturbations reveals that not only the involved cell ensembles, but also the temporal changes in these ensembles contain important, stimulus specific information. Perturbations between the repeats (R/R′) had only a small, although not negligible, effect on predictability, which suggests that either the correlative information between cell identities exists, but it does not play a major role in the encoding, or the datasets were not robust enough to reliably extract this kind of information. The results of the perturbation analysis for the LDA method were very similar (Fig. S3), except for its consistently low performance for the order-encoded dataset, confirming the above arguments.

Our results are in line with the overall picture that when a sound is heard, the auditory cortex first responds with transient discharges across a larger population of neurons. Then the activation becomes gradually restricted to a smaller population of neurons, which results in a selective representation of the sound across both neuronal population and time [32]. Higher level features, like frequency modulation or species-specific vocalizations, can be extracted by non-tone-responsive neurons, which are selective for more complex stimuli, and their responses are also often context-dependent [33]. Wang [32] argues that the combination selectivity is a general organizational principle for cortical neurons across many species. According to Smith and Lewicki [34], a complete acoustic waveform can be represented efficiently with a population spike code, that could provide a solid base for the extraction of higher features.

From a methodological point of view, our results illustrate that sequence-based neural networks, like recurrent networks or transformer networks [30], are promising tools as neural decoders. Their main advantage, compared to classical decoder models, is that we do not need manual feature extraction, because the feature extraction is performed by the network itself. It also means that, instead of a feature vector (like population vectors), the input of the model can be the raw spike train data. This is a useful property if we are unsure of the involved features. The results for the experimental datasets show the presence of population encoding for auditory stimuli. The RNN and LDA performed similarly in this case. Note, however, that the latter required an extended population vector that expresses also temporal changes. But the RNN can also reveal other kind of encodings, like rank order encoding, which are challenging for the classic linear methods. The illustrated use of random references can be a useful amendment in similar studies, which enables the testing of specific coding hypotheses.

## 4 Conclusions

Neural network-based models need a large amount of high quality data. Recent advancements in micro-electrode techniques can provide the necessary temporal precision, and the number of recordable cells also increases rapidly. However the integration of individual-specific recordings is still problematic, because there is no mutual correspondence between the single-cell recordings of the individual experiments. Consequently, cell-population based decoding models need to be fitted separately. The combination of microscopic and microelectrode techniques could help to remedy this shortcoming (see e.g. [35]); the parallel recording of morphological or spatial information would enable the classification of cells on a common positional, morfological or developmental ground, opening new possibilities also for the analytical mehods [36,37].

## Acknowledgements

This work was supported by the GINOP-2.3.2-15-2016-00057 research grant, the Hungarian Scientific Research Fund OTKA K119650, the National Brain Research Program 2017 1.2.1-NKP-2017-00002, the National Research, Development and Innovation Office (NKFIH) under Élvonal project (“Evolution and Learning”) grant number KKP129848, and by the Templeton Foundation Diverse Intelligence Grant under grant number TWCF0268 (“Learning in evolution, evolution in learning”). All the datasets and the RNN decoder code employed in this study will be available online upon acceptance.

## Supplementary Information

**Fig. S1.**
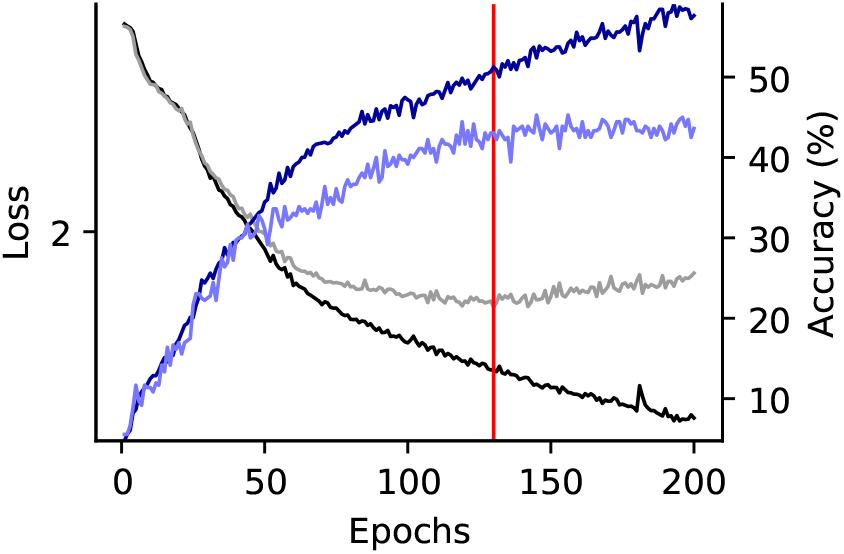
RNN decoder training session for the D^E^_I_ dataset. Black and gray lines show the loss values, while dark and light blue lines show the accuracies for the training and evaluation subsets, respectively. The red line shows the point where the evaluation loss reached its minimum and the model was saved.

**Fig. S2.**
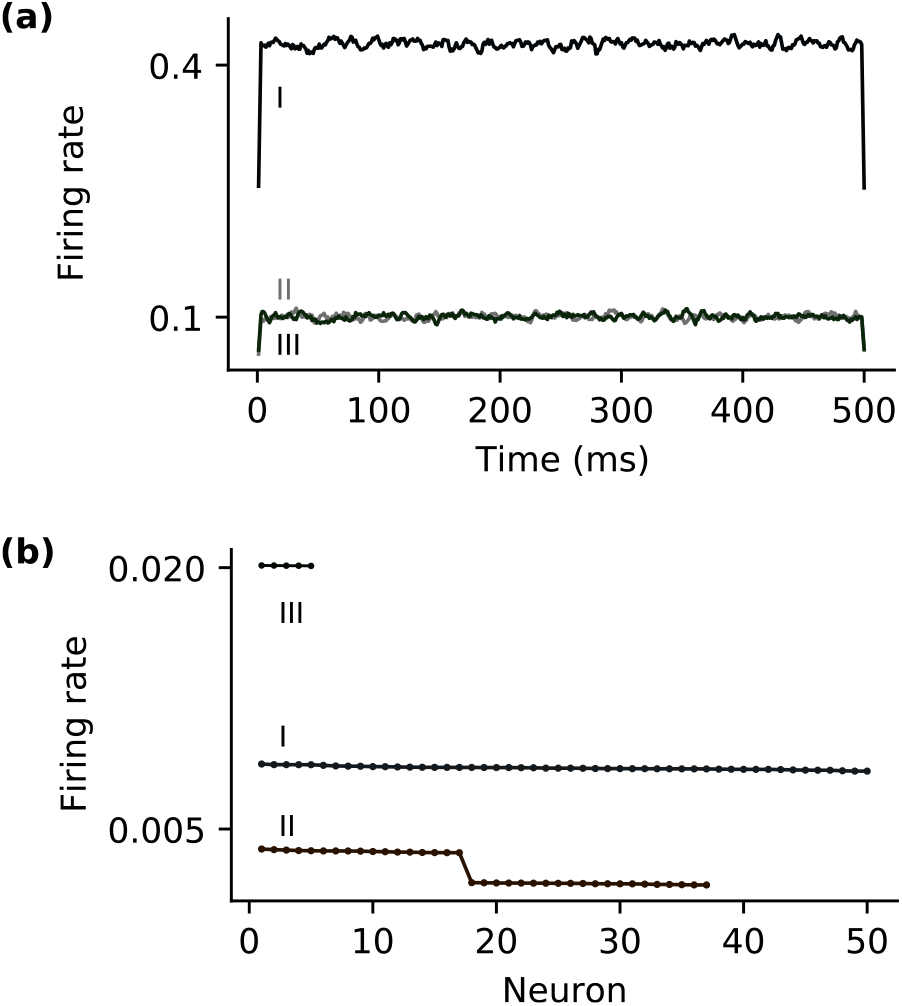
Descriptive statistics of the timebin- and cell-specific firing rates in the artificial datasets. **a** Overall firing rates, summed over all cells, in subsequent time bins after the stimulus onset. The timeseries were smoothed with a flat window, five bins wide. **b** Overall firing rates, averaged over time bins, of different neurons in decreasing order.

**Fig. S3.**
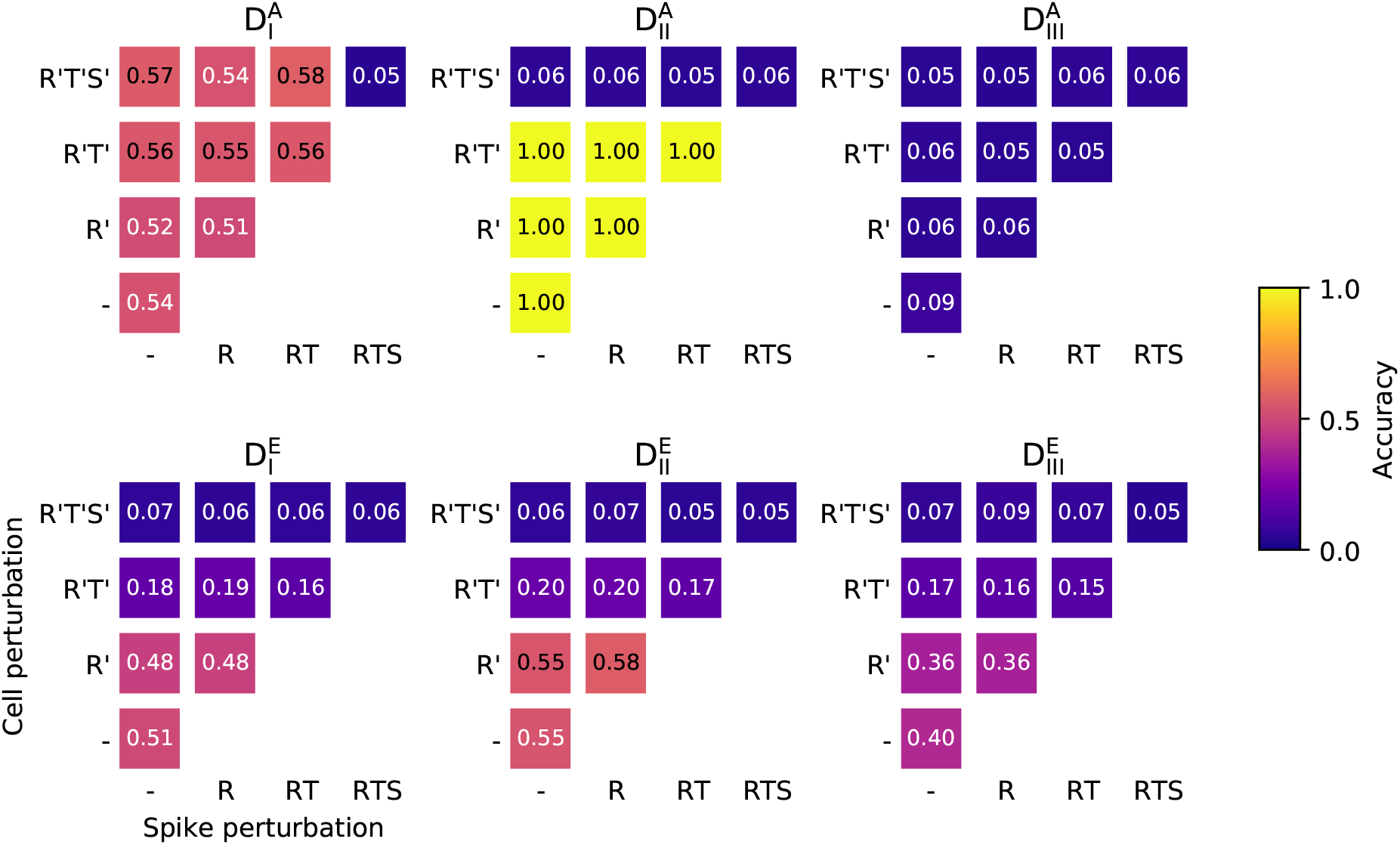
Perturbation analysis for the LDA decoder. The prediction accuracy of the LDA decoder for the original datasets is compared with the prediction accuracies for the perturbed datasets. Perturbations may affect either any of the values (spike perturbations) or only the positive values (cell perturbations) within the datasets. R, RT, and RTS spike perturbation names (and their primed counterparts for cell perturbations) refer to the affected repeat, time, and stimulus axis combinations (see main text for details).

